# Latent-space embedding of expression data identifies gene signatures from sputum samples of asthmatic patients

**DOI:** 10.1101/646976

**Authors:** Shaoke Lou, Tianxiao Li, Daniel Spakowicz, Geoffrey Lowell Chupp, Mark Gerstein

## Abstract

The pathogenesis of asthma is a complex process involving multiple genes and pathways. Identifying biomarkers from asthma datasets, especially those that include heterogeneous subpopulations, is challenging. In this work, we developed a framework that incorporates a denoising autoencoder and a supervised learning approach to identify gene signatures related to asthma severity. The autoencoder embeds high-dimensional gene expression data into a lower-dimensional latent space in an unsupervised fashion, enabling us to extract distinguishing features from gene expression data. We found that the weights on hidden units in this latent space correlate well with previously defined and clinically relevant clusters of patients. Moreover, pathway analysis based on each gene’s contribution to the hidden units showed significant enrichment in known asthma-related pathways. We then used genes that contribute most to the hidden units to develop a secondary supervised classifier (based on random forest) for directly predicting asthma severity. The random-forest importance metric from this classifier identified a signature based on 50 key genes, which can predict severity with an AUROC of 0.81 and thus have potential as diagnostic biomarkers. Furthermore, the key genes could also be used for successfully estimating, via support-vector-machine regression, the FEV1/FVC ratios across patients, achieving pre- and post-treatment correlations of 0.56 and 0.65, respectively (between predicted and observed values). The 50 biomarker candidate genes can be found in supplementary. The source codes are freely available upon request.

## Introduction

Asthma is a common chronic disease of the airways. According to a medical expenditure survey in the United States from 2008 to 2013, asthma has a prevalence of 4.8% and imposes significant economic burden, including costs due to missed work and school, medication and mortality [1].

Asthma is recognized as a heterogeneous and complex disease involving many biological pathways [2]. Among asthma patients identified as severe, subpopulations with diverse pathogenicity may exist that respond differently to medications [3]. Thus, identifying distinct subgroups of asthma patients is crucial for personalized medical decisions and management. Several researchers have investigated aspects of asthma heterogeneity and tried to identify subgroups based on different types of indicators. Simpson et al. categorized asthma patients into four subtypes, eosinophilic, neutrophilic, mixed granulocytic and paucigranulocytic asthma [4], based on the count and components of white blood cells in induced sputum [5]. The Severe Asthma Research Program performed hierarchical clustering on phenotypic measures of 856 patients and revealed five groups with distinct phenotypic features [6]. Yan et al. identified three transcriptomic endotypes of asthma (TEAs) using unsupervised clustering on gene expression of induced sputum of asthma patients, demonstrating the predictive potential of molecular profiles on disease phenotypes [7]. Each of these studies tried to interpret the identified subgroups by investigating how they associated with disease phenotypes, but did not explicitly evaluate their association with disease severity. Thus, we have yet to characterize a stringent set of indicators of severe phenotypes. As many asthma subgroups contain a non-trivial proportion of severe patients, we need to further characterize specific genes or pathways that lead to more severe phenotypes within each patient subgroup.

Scientists have extensively used transcriptome profiling to study human diseases at a molecular level. Gene transcripts that show significant differential expression and structural aberrations associated with disease phenotypes provide promising markers of clinical significance [8]. Easily obtained non-invasive biospecimens are useful sources of markers with high potential for convenient and efficient clinical applications [7]. Hekking et al. identified differentially expressed gene and pathway signatures for adult- and childhood-onset severe asthma from transcriptomes of brushings and sputum [9].

The pathogenicity of asthma involves complex and non-linear interactions between several biological pathways [10]. Thus, higher-order, non-linear features will be necessary to capture the intrinsic structure of gene expression data. We can apply non-linear generative models in order to obtain more stable representations of the data for robust feature extraction. For example, a denoising autoencoder attempts to reconstruct the original data from a randomly corrupted input, and the resulting model can potentially map the high-dimensional input data to lower-dimensional representations that are robust to small noise in the input [11]. This framework is therefore useful for extracting useful features in noisy and high-dimensional transcriptome data. Tan et al. applied the method to breast cancer gene expression data and identified features that are related to prognosis of patient survival [12].

To reveal the intrinsic structure and extract predictive features from heterogeneous transcriptome data, we propose a dAsthma framework that uses a denoising autoencoder to generate robust and non-linear representations with clinical relevance. We argue that (1) this simple structure could retrieve biological relevance and explainable components, (2) the hidden units could produce clearer patterns than the raw data to categorize patients into heterogeneous groups, and (3) the components of the clinically relevant hidden units may contain genes that are functionally associated with the pathogenesis of asthma, serving as potential sources for biomarker discovery.

## Results

### Training of the denoising autoencoder

We used a one-layer denoising autoencoder model comprised of an encoder and a decoder (Fig 1). The encoder embeds the original input data into a lower-dimensional space, the hidden layer, and the decoder reconstructs the input from the values of the hidden layer. We tuned the parameters of the model using cross-validation by training on 70% of the sample and testing on the left-out 30%. Both the training and testing loss showed proper convergence (Fig S1), indicating that we largely avoided overfitting. We then projected the input data from all non-control input samples to the embedding space of the trained model, obtaining a set of 50-dimensional vectors.

**Fig. 1.**
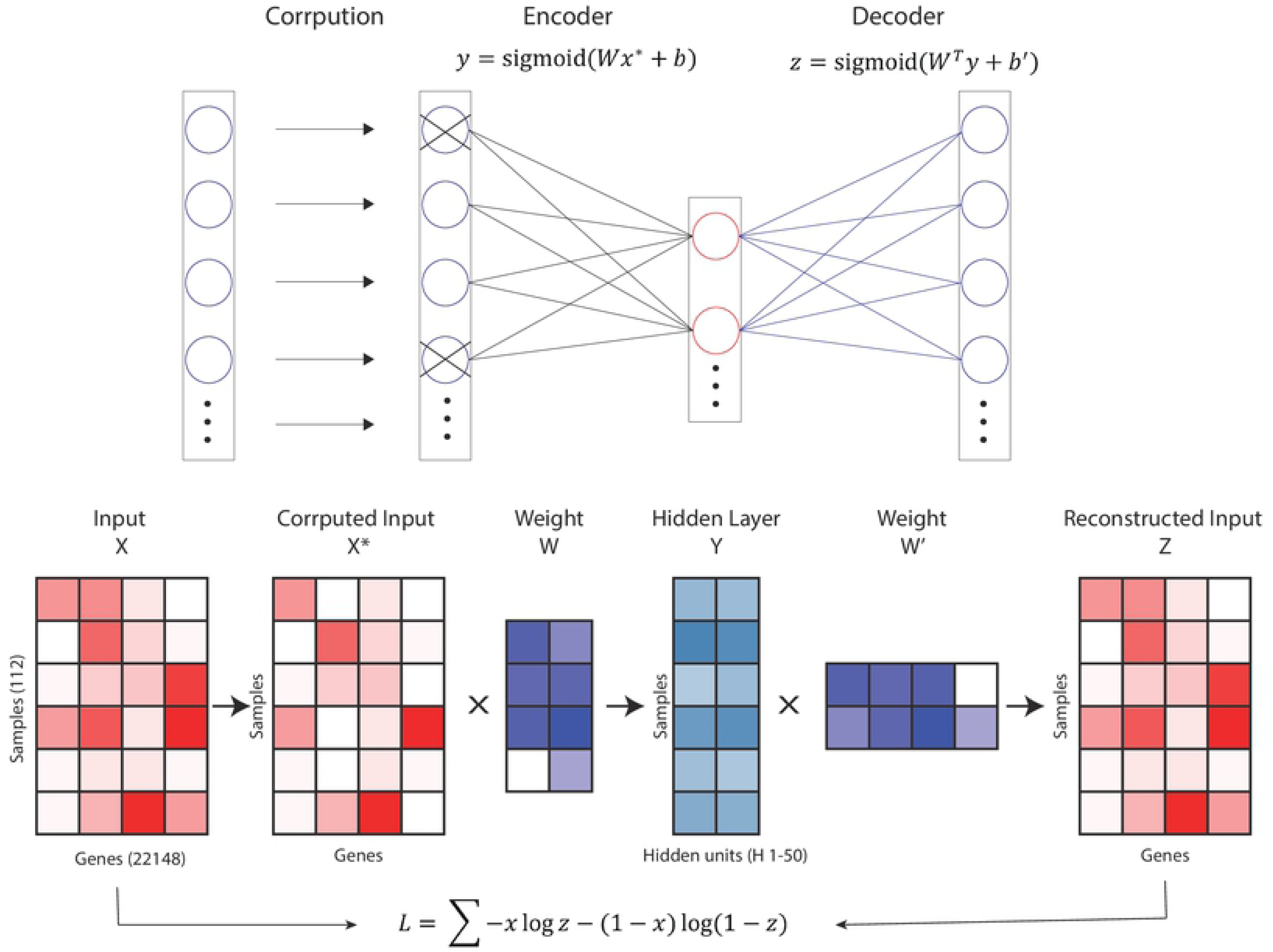
The denoising autoencoder model architecture.

### Hidden units associate with TEA clusters

By encoding the original data into the hidden vector space, the model produced a sparse embedding space; moreover, we could observe distinct patterns related to clinical traits (Fig 2a). Some hidden units showed approximately complementary behaviors (e.g., H26 and H38), indicating their distinct relevance to key molecular mechanisms and clinical subgroups of patients.

**Fig. 2.**
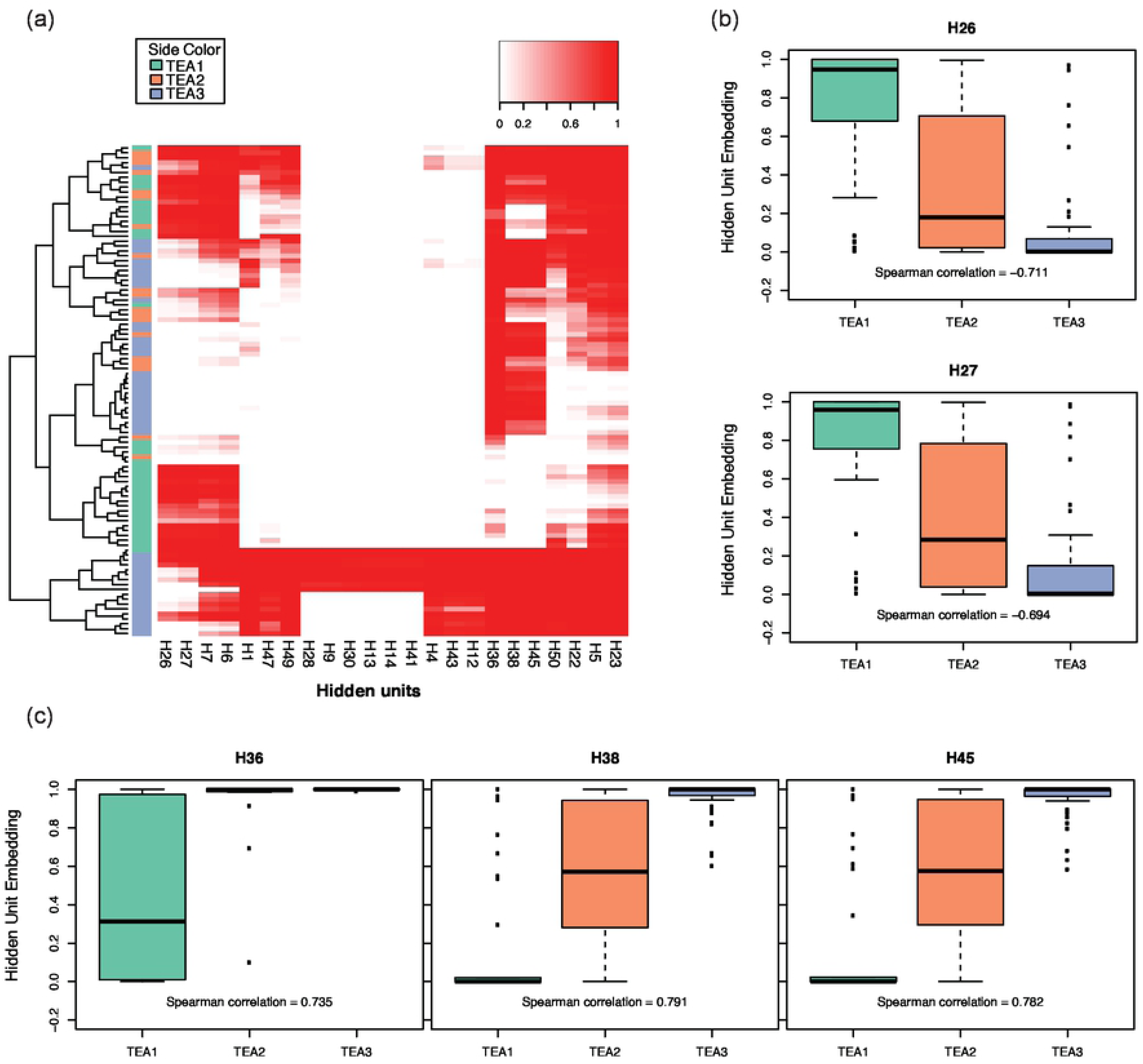
Identification of clinically-relevant hidden units. **(a)** Heatmap of hidden layer embeddings. The sidebar indicates the TEA cluster assignment of the sample. **(b)** Hidden units that are negatively correlated with the TEA cluster label. (H26, H27) **(c)** Hidden units that are positively correlated with the TEA cluster label. (H36, H38, H45)

To better interpret the learned patterns, we evaluated the embeddings of all input data for their correlations with identified clinical traits. We first removed hidden units with variance < 0.001. Using the embeddings of the highly-variable hidden units, the samples formed several small clusters corresponding to the previously defined TEA clusters, with high homogeneity (Fig 2a). It was found that TEA1 containing larger proportion of severe patients than TEA3 (Fig S2). TEA2 was relatively harder to distinguish because it was somewhat of an intermediate between TEA1 and TEA3, as some of the TEA2 samples were spread across several clusters.

Generally, the values of hidden units showed gradual monotonic changes, either increasing or decreasing, from TEA1 to TEA3 (Fig S3), indicating associations of the hidden units with clinical traits of the samples. We selected five hidden units (H26, H27, H36, H38, H45) that were significantly correlated with TEA cluster labels (Spearman correlation > 0.65), namely *Hsig*. Based on the performance of these five hidden units, we could further categorize them into two major classes: one that was negatively correlated with TEA cluster labels (H26, H27; Fig 3b), and one that was positively correlated (H36, H38, H45; Fig 3c).

**Fig. 3.**
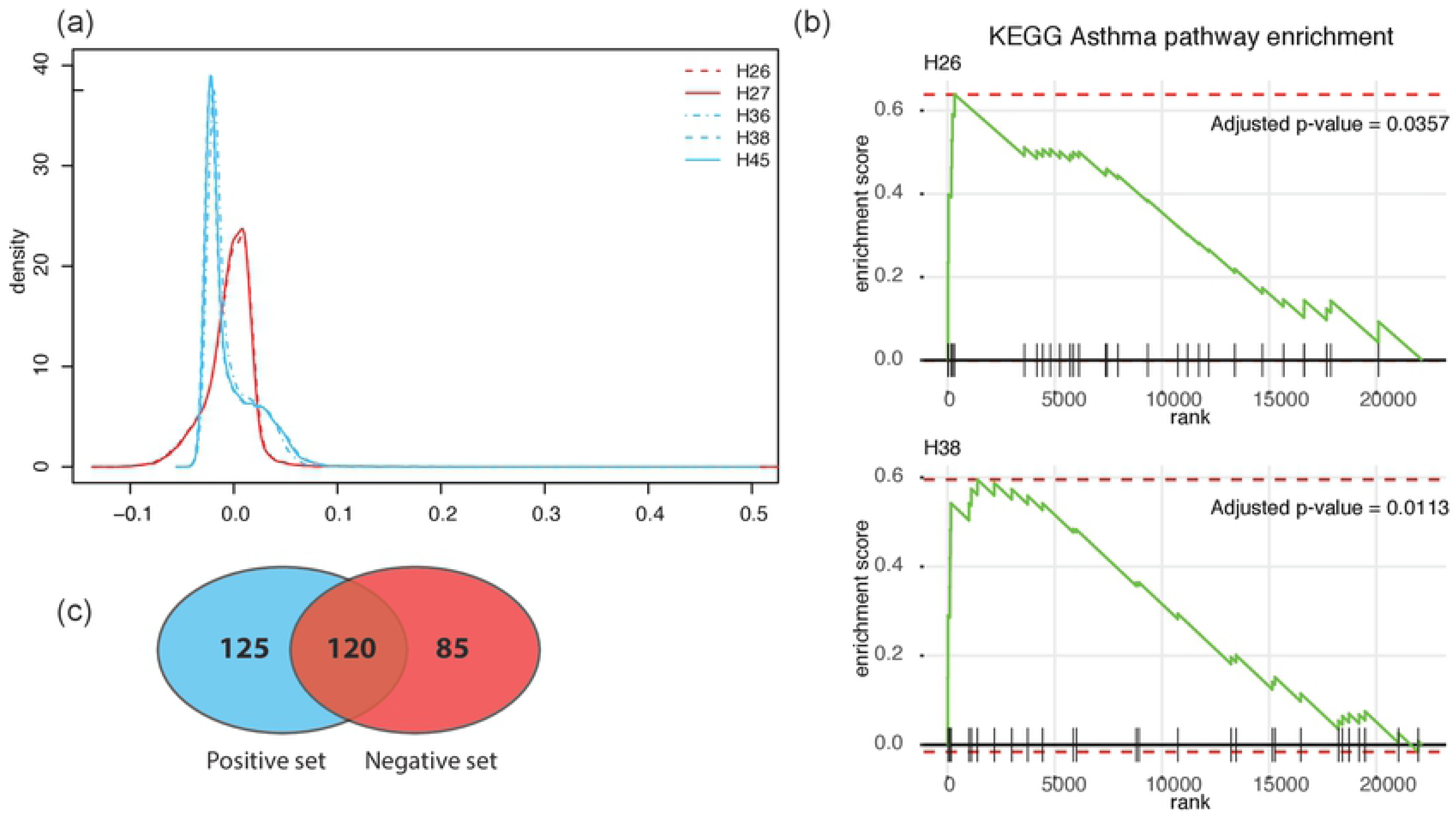
Annotation of hidden units. **(a)** Distribution of encoder weights of *Hsig*. The positive and negative sets show very different distribution patterns. **(b)** Gene set enrichment of asthma pathway from KEGG (KEGG_ASTHMA) for H27, H36 and H45. **(c)** Heatmap of encoder weights for top-weighted genes. The sidebar indicates whether the gene uniquely belongs to the negative set or positive set.

### Annotation of relevant hidden units

To further understand the biological significance of the hidden units, we tried to associate them with functional enrichment based on the weights of the encoder network that mapped the input to the hidden unit space. This weight represents the contribution of the gene to the value of the hidden unit, and can be considered the component of the hidden unit. Thus, we could infer biological significance of the hidden units from the weights of the encoder.

The weight distribution of the encoder layer showed similar patterns for hidden units belonging to the same set (Fig 3a). Comparatively, the negative set showed a nearly symmetric distribution with a slight negative skew, whereas the positive set showed a highly positively-skewed shape.

Using encoder weights for all 22,148 genes as the ranking score, we performed gene set enrichment analysis using KEGG pathway gene sets to obtain functional enrichments of *Hsig*. Similar to the weight distribution, hidden units from the same set showed strong resemblance with respect to enrichment of functional terms (Fig S4). Notably, many of the enriched functional terms were related to molecular functions and signaling pathways associated with disease pathogeny and autoimmune response, whereas deficient terms included “gene expression machinery” and “metabolism”. Specifically, the enrichment of the term “asthma” showed high significance in all five hidden units (Fig 3b, Fig S5).

We then extracted the top-weighted genes of *Hsig* as potential clinical markers for downstream analysis. For each hidden unit, we extracted the genes with top 200 highest weights; we removed ribosome-related genes from the analysis to prevent ribosome functional roles from dominating the selected gene set. Concordant with previous observations, the gene sets for hidden units belonging to the same class were highly similar. By merging the top-weighted genes from all five hidden units, we obtained a list of 330 genes. We identified several genes that showed distinct differential patterns of weights between the positive and negative set (Fig 3c). GO analysis revealed an enrichment of disease-related terms like “antigen processing and presentation”, “immune response” and “interferon-gamma signaling pathway” (Fig S6).

### Prediction of asthma severity

In order to assess the clinical relevance of the learned model, we investigated the association between hidden unit values and clinical data. Generally, the performance of hidden units showed distinct correlations with various clinical traits (Fig S7). We then tested the predictive performance of some clinical traits using the embedding values of *Hsig* and the expression of the combined list of their top-weighted genes.

We first used the value of all hidden units, *Hsig* and the top-weighted genes to predict the asthma severity levels of the patients. We only used samples labeled as “mild” or “severe” for the classification analysis. Given each training dataset, we trained a random forest classifier and assessed the predictive accuracy on the test data, represented by the AUROC value (Fig 4a).

**Fig. 4.**
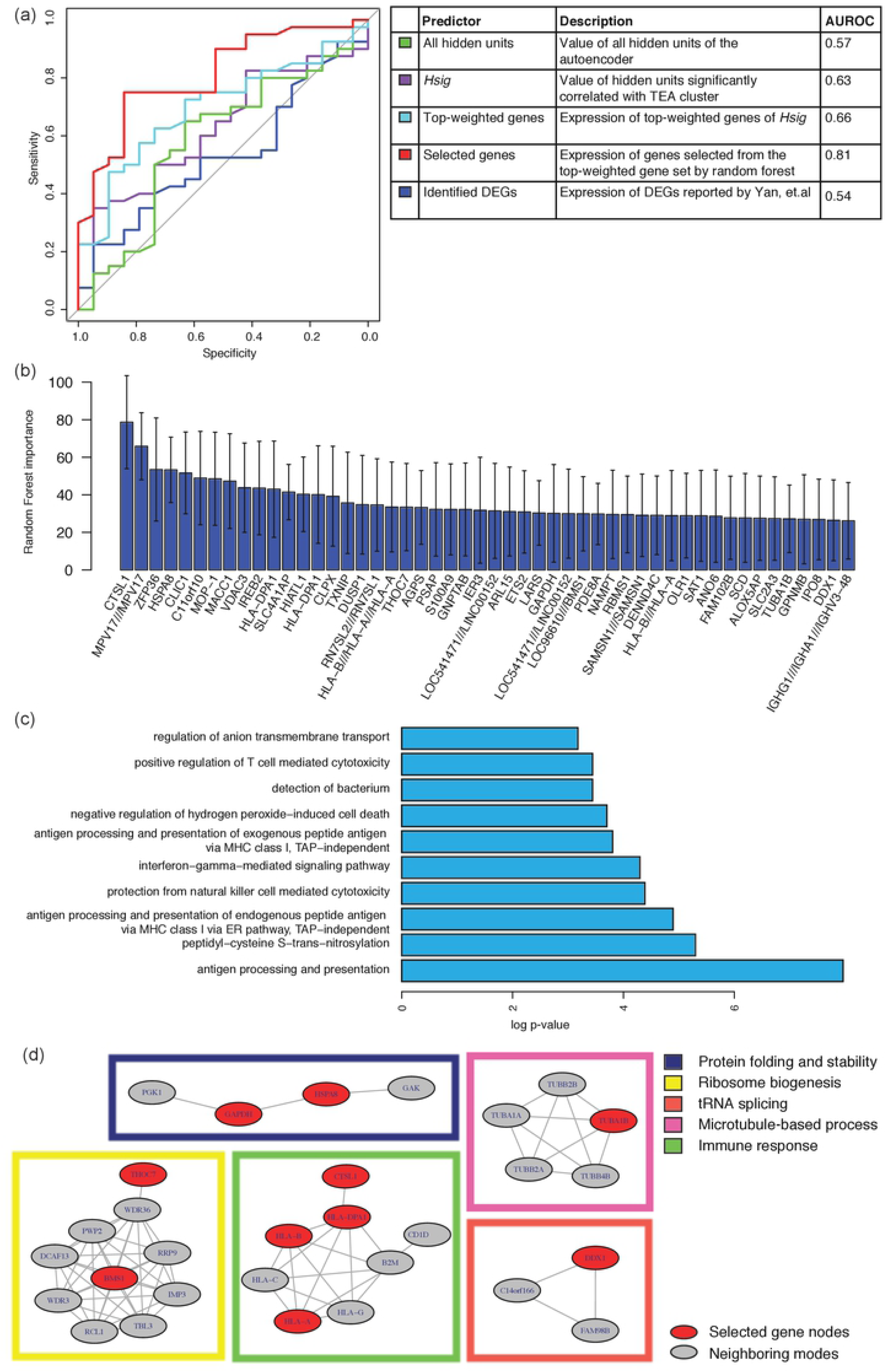
Prediction of asthma severity and feature selection. **(a)** AUROC of the prediction of asthma severity using different feature sets. **(b)** Importance of the selected genes. **(c)** GO term enrichment of the selected genes. **(d)** Selected genes in the context of PPI network.

From the top-weighted genes, we then performed feature selection to obtain the most relevant genes. We considered 50 genes with the top average importance as the most relevant to the prediction of severity (Fig 4b, Table S1). Expression of these genes reached an average AUROC of 0.8066 (0.6194 for random selection of 50 genes from top-weighted gene list) in predicting severity level. Comparatively, this value is higher than for a set of differentially expressed genes between asthma patients and control samples identified by Yan et al. (average AUROC of 0.5448; Fig 4a). We hypothesize that asthma-associated genes, identified from case-control differential expression analysis, may represent a collective set of genes in general pathogenic pathways. As pathogenicity and transition to severe phenotypes may involve different mechanisms, they may not be sufficient to fully explain the causes of severe asthma for different subgroups.

Our list of selected genes included genes related to autoimmune responses, such as antigen processing and presentation, T-cell toxicity, interferon signaling and cell adhesion (Fig 4c). In the context of a protein-protein interaction network, we were able to identify genes residing in various functional modules (Fig 4d). Specifically, several genes, such as human leukocyte antigen genes and cathepsin L1, belonged to a module related to immune response. In addition, the list included genes with potential relevance to asthma pathogeny, such as immunoglobulin heavy chain genes (IGHG1, IGHA1 and IGHV3-48), inflammation (S100A9) [14] and iron response (IREB2) [15, 16].

### Prediction of clinical measurements

To evaluate an individual’s lung function, clinicians use the FEV1/FVC ratio, which is the ratio of the volume forcefully exhaled in a second versus the maximum volume of a forceful exhale [17]. A significantly reduced FEV1/FVC ratio is a sign of airflow obstruction, and is considered a criterion for asthma severity.

We found that the hidden units in *Hsig* generally showed a higher absolute value of correlation with the FEV1/FVC ratio, in both the positively and negatively correlated subsets, compared to other hidden units, indicating clinical relevance of these hidden units (Fig 5a).

**Fig. 5.**
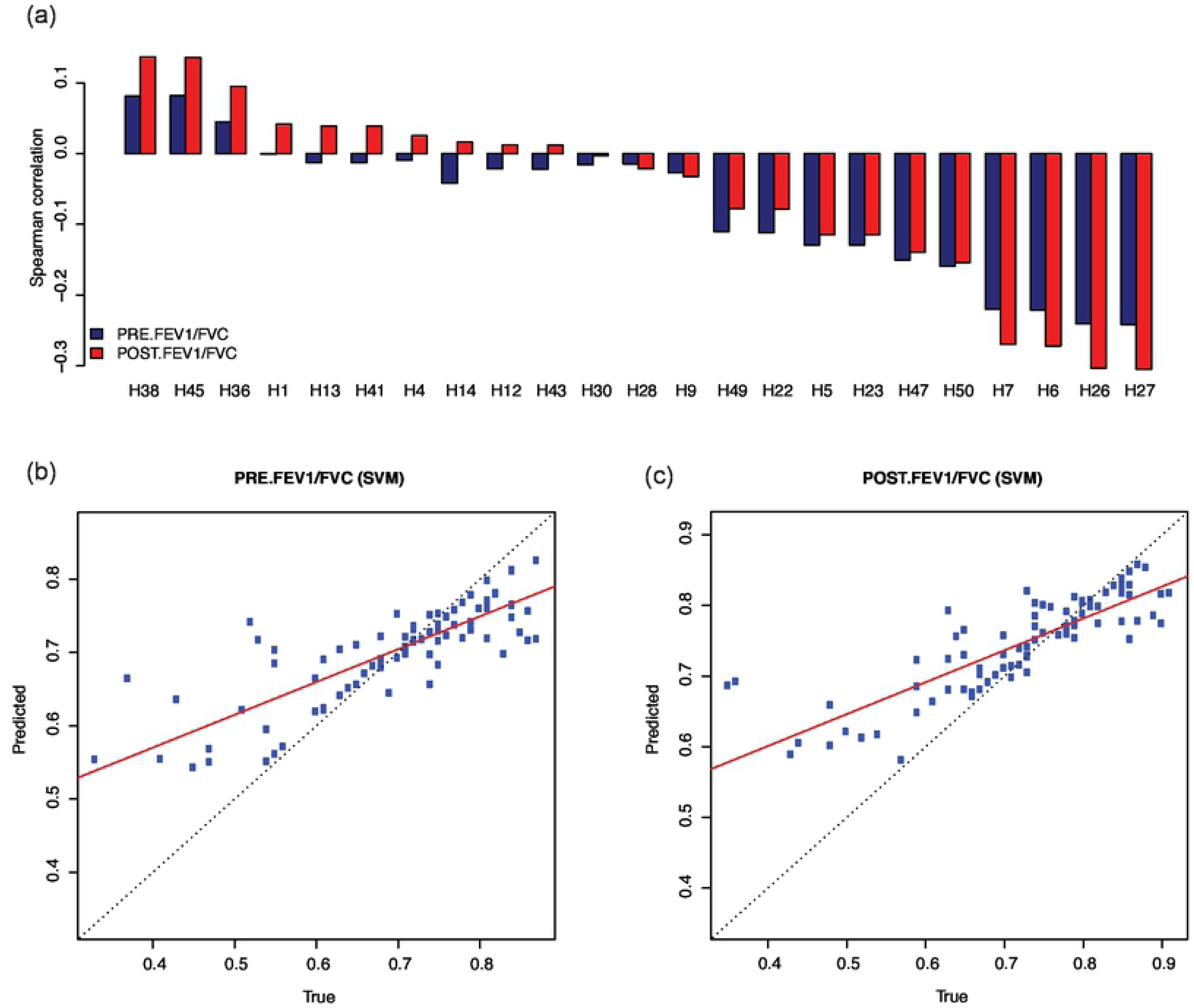
Prediction of FEV1/FVC ratio. **(a)** Spearman correlation between hidden unit values and pre-/post- treatment FEV1/FVC. **(b)** Plot of predicted value versus true value of pre-treatment FEV1/FVC using support vector machine regression with selected gene expression. **(c)** Plot of predicted value versus true value of post-treatment FEV1/FVC using support vector machine regression with selected gene expression.

Finally, we tried to predict the pre- and post-treatment FEV1/FVC ratios with the value of hidden units and expression of their top-weighted genes using a support vector machine and LASSO (Fig 5b, c, Table S2, S3 and Fig S8, S9). The selected genes achieved the highest predictive performance, in terms of both mean squared error and explained variance. Together, these results show that the selected hidden units and genes are clinically relevant and could be used as markers for asthma severity.

## Discussion

Research suggests that asthma is a heterogeneous disease, as its pathogenicity may largely depend on a patient’s individual genetic variability [18]. For several previously-proposed subsetting methods of the asthma population, many of the sub-populations contains non-trivial proportion of severe cases [7]. This indicates that severe asthma may be the result of multiple subtype-specific mechanisms. Thus, diagnosis from disease phenotypes solely may not provide sufficient information for personalized treatment. To identify the molecular regulatory mechanisms associated with asthma severity, we developed a framework called dAsthma using a denoising autoencoder model trained on gene expression profiles of sputum from asthma patients. The model can map high-dimensional expression data to a lower-dimensional latent vector space composed of 50 hidden units, and cluster the patients into clinically relevant subgroups using the embeddings in the latent space. We then investigated how the hidden units were associated with biological pathways and clinical traits. From the most relevant hidden units, we discovered a set of 50 genes whose expression profiles combined could predict asthma severity with high accuracy.

The dAsthma framework learns a more robust representation of the data by adding random noise to the input data. It looks for definitive features that account for the variations among the dataset. Compared with raw gene expression data, which produces less distinguishable patterns in clustering analysis, the value of the hidden units learned by the dAsthma model can identify distinct clusters out of the noisy data. We note that these clusters are generated in an unsupervised fashion, that is from mere gene expression data with no external information provided.

We tuned the autoencoder model of our framework to make it more robust. We selected an optimal learning and corruption rate by performing a grid search of hyper-parameters. The dataset that we used as training set for our model was relatively small; this may limit the ability of the autoencoder to identify underlying patterns of the data due to lack of information, and could introduce additional risk of overfitting. We tested our model using cross-validation and found good convergence of both training and test loss. We selected the model architecture, that is the number of hidden nodes, based on following considerations. As smaller number of hidden units may fail to capture some subtle structures among the dataset, we used a relatively larger and presumably redundant number (50 hidden units), so that we could retain as much useful information as possible. We then filtered the 50 learned hidden units for better conciseness and specificity. Among these 50 hidden units, some showed almost uniform values across all samples; these were of less interest and were discarded. In addition, some highly variable hidden units had similar performances and components. We collapsed these highly correlated hidden units for downstream analysis.

Neural network-like models become less interpretable as they grow deeper and more complex. Especially when applied to biological data, associating a learned model with underlying mechanisms of molecular functions and disease pathology is challenging. Our dAsthma framework uses a simple, low-complexity structure to achieve better interpretability. The learned patterns can be interpreted from two perspectives. The first is to study the enrichment of biological pathways of the hidden units based on their components (i.e., genes weighted by their contributions to the hidden units). The second is to associate the patterns with external information about clinical measurements, such as asthma severity. These analyses show that the hidden representations learned by the denoising autoencoder could bridge gene expression and clinical traits. Overall, the hidden units can be recognized as “gene modules” that represent some key biological pathways in the pathogenicity of asthma, which leads to various disease phenotypes.

The definition of asthma severity is mainly based on phenotypic traits, and may vary across studies. The 50 severity-associated genes selected from the components of the most clinically relevant hidden units potentially could be used to characterize asthma severity for different subtypes, as the denoising autoencoder in dAsthma tends to identify features that account for variations among the asthma patient population.

To summarize, we have shown the strength of our dAsthma framework, which makes use of a denoising autoencoder and produces biologically interpretable results, to extract clinically relevant patterns and select potential biomarker genes from gene expression profiles of sputum of asthma patients.

## Materials and Methods

### Data

The raw expression data is provided by Yan [7], After quantile normalization, all expression data is scaled to [0,1] by min-max method.

### Implementation and training of the denoising autoencoder

The denoising autoencoder is comprised of two symmetric neural networks: an encoder network and a decoder network. The encoder network first maps a corrupted input *X* *, i.e. original input data *X* with randomly zeroed-out features, to a hidden vector space *Y*:

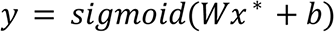

The decoder network then tries to produce a reconstructed input *Z* from the hidden vector space that resembles the original input as much as possible:

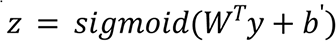

Specifically, we constrain the weights of the decoder network to be the transpose of that of the encoder. The loss function is cross-entropy loss:

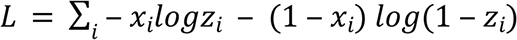

Then the derivative can be calculated as follows:

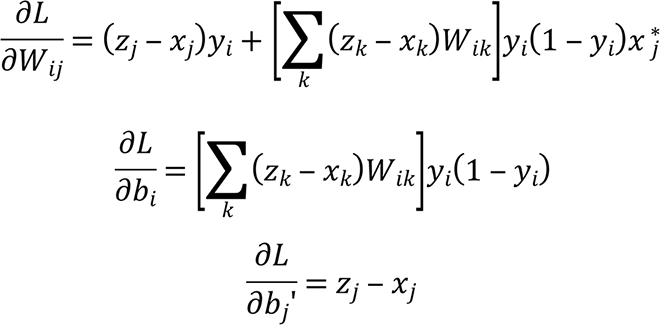

The model is implemented with C++. Specifically, weights are shared between the encoder and the decoder, i.e. the weight matrix of the decoder is a transposition of that of the encoder. The output layers of the encoder and decoder are activated with sigmoid function.

We used a model with 50 hidden units. After hyper-parameter tuning, the model is then trained with 100 epochs, with learning rate=0.1, corruption rate=0.001, to minimize the cross-entropy loss. Cross validation was performed with 7:3 random training-test data split of the original data.

### Gene set enrichment analysis (GSEA)

We use the fgsea function (from R package fgsea) [13] to perform gene set enrichment analysis. The fgsea function expects an input of statistic array for genes in the gene list of interest, as a measurement of the relevance of the genes with the desired phenotype. In our scenario, for each hidden unit, we regard the learned weights of the input features (genes) as the statistic array. We use the curated KEGG gene set from MSigDB (downloaded from http://software.broadinstitute.org/gsea/downloads.jsp) for the analysis. The output is visualized as the plot of enrichment scores against genes ranked by the statistic values.

### Prediction of clinical traits

Generally, prediction of severity, pre- and post-treatment FEV1/FVC ratio are evaluated using 4-fold cross validation. The data is randomly split into four equal parts. For each part, the target value is predicted using a model trained with the other three parts. The predicted values of the 4 parts are then concatenated and compared to the true values.

For prediction of severity, only samples labeled as “Mild” and “Severe” are used for the analysis. We perform random forest (using R package caret) on the training data with default parameter settings. Alteration of parameters, i.e. the number of trees and number of features for splitting of the nodes, does not significantly impact the results (data not shown). The AUROC reported is the average over 10 repeats of 4-fold cross validation.

For prediction of pre- and post-treatment FEV1/FVC ratio, both support vector machine (SVM) regression (using R package e1071) and lasso (using R package glmnet) are tested. The predictive power of the trained model is assessed by calculating the Pearson correlation between predicted values and true values.

### Feature selection

A initial gene set is generated by merging the top-weighted genes for the five most clinically relevant hidden units. The merged gene list, containing 330 genes, are used as input to a random forest classifier to predict the severity label (“Mild” or “Severe”), with all samples used. For a random forest regression model, the importance of input variables is calculated as follows: for each tree, the out-of-bag MSE is computed before and after randomly permuting a variable. The importance of the variable is defined as the average difference of the out-of-bag MSE before/after permutation over all trees.

The importance of the features in the learned model is then extracted from the learned model. We use 50 genes with highest importance averaged over 50 trials as the selected genes for downstream analysis.

### Network analysis

Proteins corresponding to the selected genes are provided to STRING (https://string-db.org/). For microarray probes targeting multiple proteins, all the corresponding proteins are included. To visualize the role of these proteins in the context of protein-protein interaction (PPI) network, we also included nodes of highly reliable first-shell interactions (colored in grey in Fig 4d) with the query proteins (colored in red in Fig 4d). Only experimental validated and database curated interactions are included.

